# Tigers in North Dakota; Tiger Salamander (*Ambystoma tigrinum*) eDNA Surveying for Presence in Cattle Ponds

**DOI:** 10.1101/2024.05.04.592111

**Authors:** Evan R Showalter

## Abstract

Environmental DNA (eDNA) allows for efficient and convenient data collection using DNA segments to confirm species presence. There is little literature using eDNA to specifically examine tiger salamander presence in North Dakota. I sampled 8 different cattle ponds in Western North Dakota twice each using a set eDNA collection method from the United States Department of Agriculture. After collecting water samples, the filters were sent to Jonah Ventures for DNA extraction and metabarcoding. Samples were incubated, extracted, ran through a first round of Polymerase Chain Reaction (PCR) inspected on agarose gel. Then the amplicons–product of an amplified or replicated piece of DNA, were cleaned, a second round of PCR followed, then the amplicons were cleaned again. Finally sequencing and bioinformatics took place. Salamanders (genus *Ambystoma*) were detected at 5 of the 8 sites. Average single season Bayesian occupancy point estimate was 0.655 ± (.165), with a detection probability of 0.735± (.156) over the two site visits. Of the covariates including elevation, surface area, and air temperature, air temperature on visit 2 had the best model but still was not significant. There was no significant correlation between any of the covariates and the naïve detections. This method was effective in detecting the genus *Ambystoma*, however more work and greater sample site variation could elucidate impacts of species level presence absence data.

## Introduction

Environmental DNA (eDNA) is an effective and efficient method for detection of elusive organisms (Harper et al., 2018; Kaganer et al., 2022). Environmental DNA or eDNA is a method of collecting data that is quickly becoming more prevalent in the landscape of sampling for presence of rare or elusive species (Hinlo et al., 2017; Ruppert, Kline & Rahman, 2019; Shirazi, Meyer & Shapiro, 2021; Schenekar, 2023). It is useful because anytime an organism leaves extracellular DNA it is creating eDNA. This could be many things such as secreted feces, mucous, or haploid gametes; shed skin and hair; and carcasses (“Environmental DNA (eDNA) | U.S. Geological Survey”). This extracellular DNA can be collected via water or soil samples making it efficient. (Harper et al., 2019; Andres et al., 2023). When collecting a water sample the individual only needs to interact with the water source and leave one of the forms of DNA mentioned behind. Once this has happened samples are collected by filtering this water. Then taking the filters and examining them through a polymerase chain reaction (PCR) which will amplify segments of the target species DNA. If DNA of the target species is found and amplified in the sample then presence of the individual of interest can be confirmed. Leading to why it can be helpful for detecting species that are cryptic. With this method long as there has already been primers found to identify the target individual, there is no need to find and detect these individuals in the field through traditional methods. They only need to interact with the area of interest and you can detect them there with fragments of their DNA that are left behind. With the assumption that the eDNA has not degraded due to temperature, pH, or UV radiation and is still able to be detected (Strickler, Fremier & Goldberg, 2015).

Tiger Salamanders (*Ambystoma tigrinum)* are cryptic and uncommonly detected in western North Dakota. I have lived in ND for 22 years spending much of my life outdoors be it hunting, fishing, or farm work and have detected the species once. They are documented in the area (North Dakota State University, North Dakota Game and Fish, HerpMapper, and North Dakota Herp Atlas). Tiger salamanders are typically 15-20 cm in length with many unique bi-color patters of spots and stripes. They may have dark gray, brown, or black bodies with brownish-yellow markings. In the United States individuals can be found along the Atlantic coast south of New York and down to Florida. With the majority living in the center of the country from Arizona and Montana east to Ohio and Kentucky (National Wildlife Federation). They live near vernal pools, ponds, and other lentic bodies of water. During winter and most the year they are burrowed underground but otherwise use many different damp areas even small mammal burrows. Some individuals will never go through metamorphosis to become land dwelling adults, retaining larval characteristics such as gills (Johnson, 2015). They still can become sexually mature and live their whole life completely underwater this condition is known as neoteny. These traits and reproductive behavior of using small bodies of water from early spring to summertime, makes them a perfect candidate for eDNA because they will be in contact with the water leaving behind forms of extracellular DNA (David Sever & Dineen, 1977; Loredo & van Vuren, 1996). Individuals should be in the area for feeding as well as reproductive opportunity (Modrow, 2008) assessed tiger salamanders in North Dakota and found exclusively aquatic prey in stomach contents of transformed salamanders. Supporting this further was that many of these transformed individuals were still juvenile and not sexually mature. EDNA samples have been collected for amphibians in cattle ponds such as (Goldberg, Strickler & Fremier, 2018). However, tiger salamanders specifically have not been assessed in cattle ponds of ND as the source of eDNA collection.

In this we assessed cattle ponds and other small ponds on properties north of Dickinson North Dakota to evaluate the hypothesis that using eDNA sampling in cattle ponds can yield positive tiger salamander detections. Being that a small lentic body of water with few predators like fish should make a suitable place for tiger salamander young to be laid and grown. As well as they should be returning to these areas for feeding opportunities. The only exception to them being undisturbed and their safety would be other organisms in the ecosystem such as cattle, deer, and birds. Detection was possible, these detections were then analyzed for occupancy and compared with co-variates such as surface area of the pond to assess factors influencing tiger salamander presence.

## Methods

### Data collection overview

There are two halves in the data collection process, collecting the in-field samples and analyzing them in the lab. For the first part to obtain a study area an area North of Dickinson North Dakota bordered by 4 roads; 28^th^ ST SW, highway 22, 23^rd^ ST SW, 101^st^ AVE SW was chosen as the study area. This area was non-randomly selected with best prospects to have access to all land due to the fact I had connections to the landowners in this area. Providing a stratified random approach with random sites inside a non-random study area (Figure 1). I initially assumed all sites were accessible and contained water to give proper basis for a random sample. Then through on ground inspection and use of OnX maps all ponds found were numbered. This process was paired with if access was available to the property’s ponds. If ponds were inaccessible they were excluded. Using the random number generator and access to these ponds, 8 sites were selected. The 8 sites were visited and sampled from 12pm – 5pm mountain time, twice, a month apart on 6-25-23 and 7-29-23. Having 8 sites and 1 revisit of each (2 total visits) was a decision made off guidance from advisors of the project.

**Figure 1.**
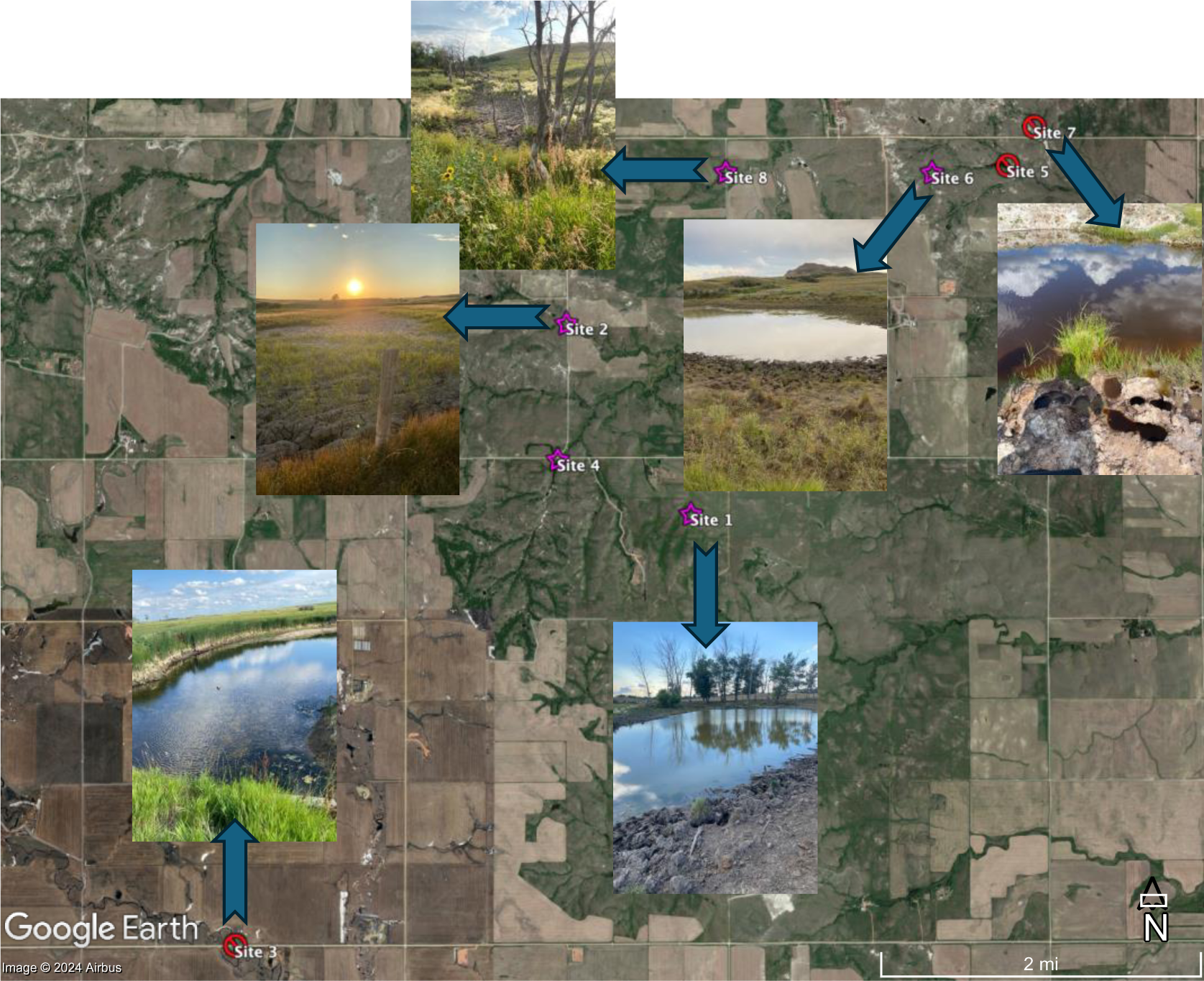
This is a map created in Google Earth Pro displaying detection or not per study site across the study area. Detections are indicated by a purple star (sites 1,2,4,6,8) and no detection is indicated by a red X (sites 3,5,7).

### In field collection Part 1: Sampling kit components

A eDNA sampling kit was used, provided by the Rocky Mountain Research Station for collection (Figure 2). This kit consisted of 3 major parts: a protected case that held a pump (2), tubing with an adapter (3), the eDNA sampling protocol (5), battery charger (13), pump battery (14), and alligator clip adapter for battery (15). An outflow bucket (8). A duffle bag containing eDNA sampling equipment (1) that held, sample box with letter sized envelopes (10), pencils, and indelible ink markers, black bag for used equipment (11), white bag with unused site kits (12). Each site kit was inside a 1 gallon resealable bag with everything you need to collect 1 sample; filter holder with filter (in clean bag) (4), forceps (in clean bag) (6), clean gloves (7), clean sample bags with desiccant (for storing filters after sample taken) (9).

**Figure 2.**
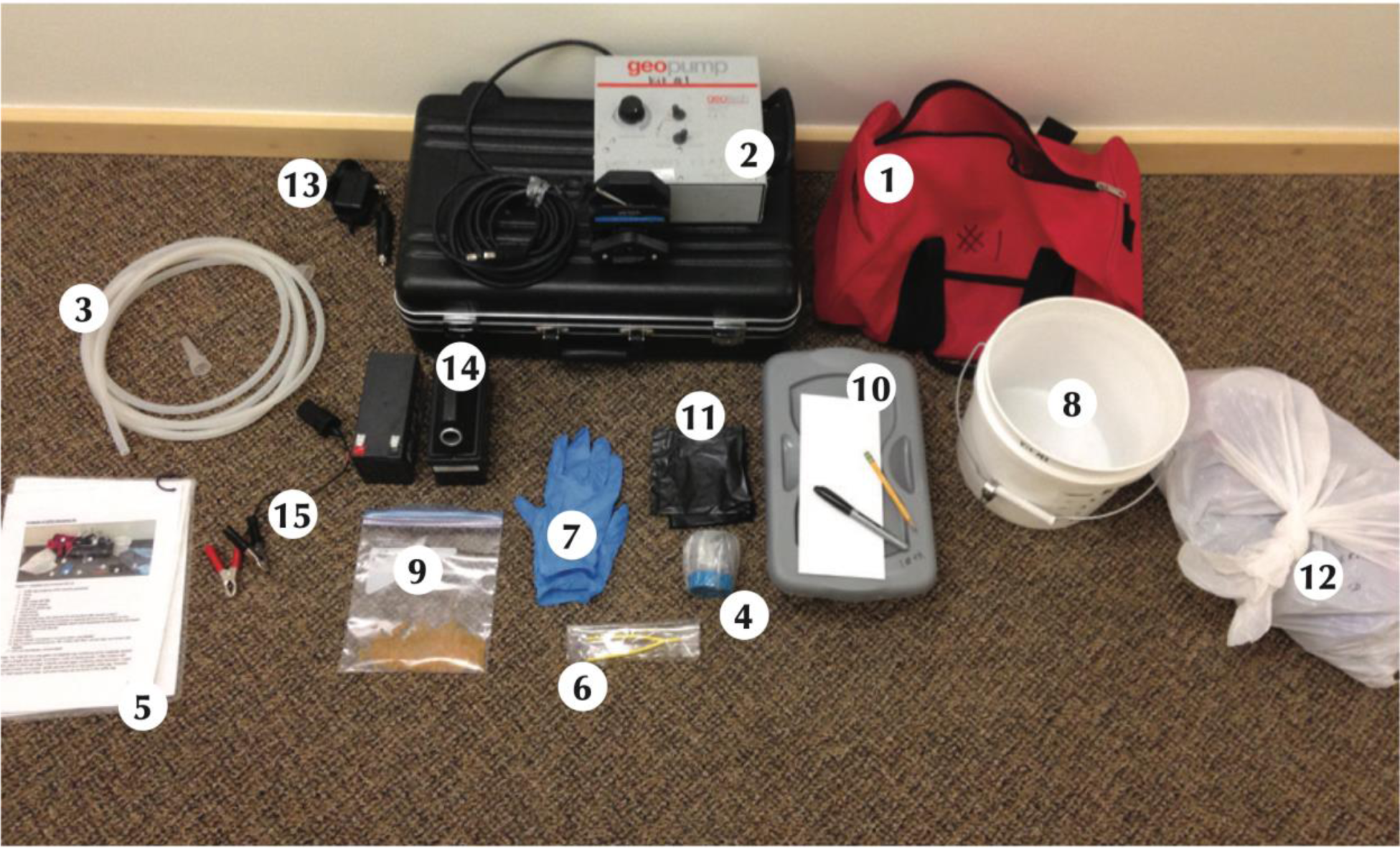
Full eDNA sampling kit with materials provided by the National Center for Wildlife and Fish Conservation. (*Photo credit: Carim et al., 2016*)

### In field collection Part 2: Sampling protocol

The Rocky Mountain Research station protocol was followed during sampling (Carim et al., 2016). **Step 1** connect the pump: removed pump, battery, and alligator clips from case and connected on a sable surface. **Step 2** install the tubing: tubing was loaded in the correct orientation as the sticker placed on the pump, for inflow and out flow ends. **Step 3** start the pump: pump started in forward direction ensuring suction. **Step 4** remove the site kit: one site kit was removed from the white bag immediately closing it after. Put on pair of gloves and made sure not to touch anything that may be contaminated with DNA. **Step 5** forceps and sample bag: removed bag containing clean forceps and the sample bag containing the silica desiccant. **Step 6** install the filter assembly: removed the packaged filter assembly from the site kit, opened but did not remove the filter from bag, held it with the bag oriented to where the tube assembly can be connected. Connected tube assembly while filter assembly was still in the bag. The hand connecting the tube was now considered the dirty hand while the hand holding the filter assembly was the clean hand. **Step 7** placement of filter assembly: submerged the filter assembly with the clean hand as now anything in the water source was part of the sample so that does not render that hand dirty. Pump runs until the water in the outflow bucket reached the marked line of 5 L. **Step 8** drain and dry the filter: once the indicated line in the outflow bucket was reached the filter assembly was pulled out of the water and the pump was allowed to run for 30 seconds drying the filter. **Step 9** remove the cup from filter assembly: at this point the cup on the filter assembly was popped off and set aside. **Step 10** fold filter and secure in sample bag: removed forceps from their protective bag. Used the forceps to fold the filter in half than in quarters with the filtrate side facing in. Placed the filter into the sample bag containing desiccant beads ensuring nothing but the filter touched the inside of the bag. Made sure the filter was at the bottom of the bag in contact with the desiccant beads and then pushed all air out and sealed the bag. **Step 11** label sample bag: gloves were removed, and bag was labeled with sample ID, coordinates, and date. **Step 12** place sample bag in envelope: an envelope was labeled with the same exact information as the bag and the bag was placed inside the envelope and sealed. **Step 13** clean up: with sample labeled and stored the rest of the sample equipment was packed up to move to the next site. All parts of the site kit were now to be placed in the black used equipment bag. **Step 14** storing samples: samples are stable for several weeks in silica desiccant beads however they should be kept away from water, heat, and sunlight. The authors recommended if its more than 2 weeks they should be stored in a freezer. These samples were stored in a -80° c freezer as that was what was available for a consistent and monitored option (Stored for approximately 25 weeks before shipped off for analysis).

### In field collection Part 3: Co-variates

The sizes of each sample location were measured with and stored in OnX maps after the second round of sampling on 7-29-23. Using the measurement tool inside the program, by selecting points around the pond a total surface area was given in number of acres. This measurement was then converted from acres to meters squared. Elevation was also collected using the measurement given by OnX maps GPS capability, it was collected in feet than converted to meters. The last covariate collected was ambient temperature at time of collection via iPhone, this was collected in °F and converted to °C. All metrics recorded, original and converted were stored in csv file and posted to open science framework (OSF) account (https://osf.io/gkz8x/).

### Lab Analysis

In the second half of the data collection the filters from each site were analyzed for detection in each site. Jonah Ventures was chosen they are a company that specializes in sequencing environmental DNA. Samples collected were shipped to Jonah Ventures where after discussion with their experts the method chose to analyze was metabarcoding with the high sensitivity assay. The best analysis that could have been done would have been a quantitative PCR. However, an assay was not available for tiger salamander, so this was the next best option that was recommended and said to be sufficient after discussion with Jonah Ventures. The metabarcoding follows a 9- step process to obtain results (Figure 3).

**Figure 3.**
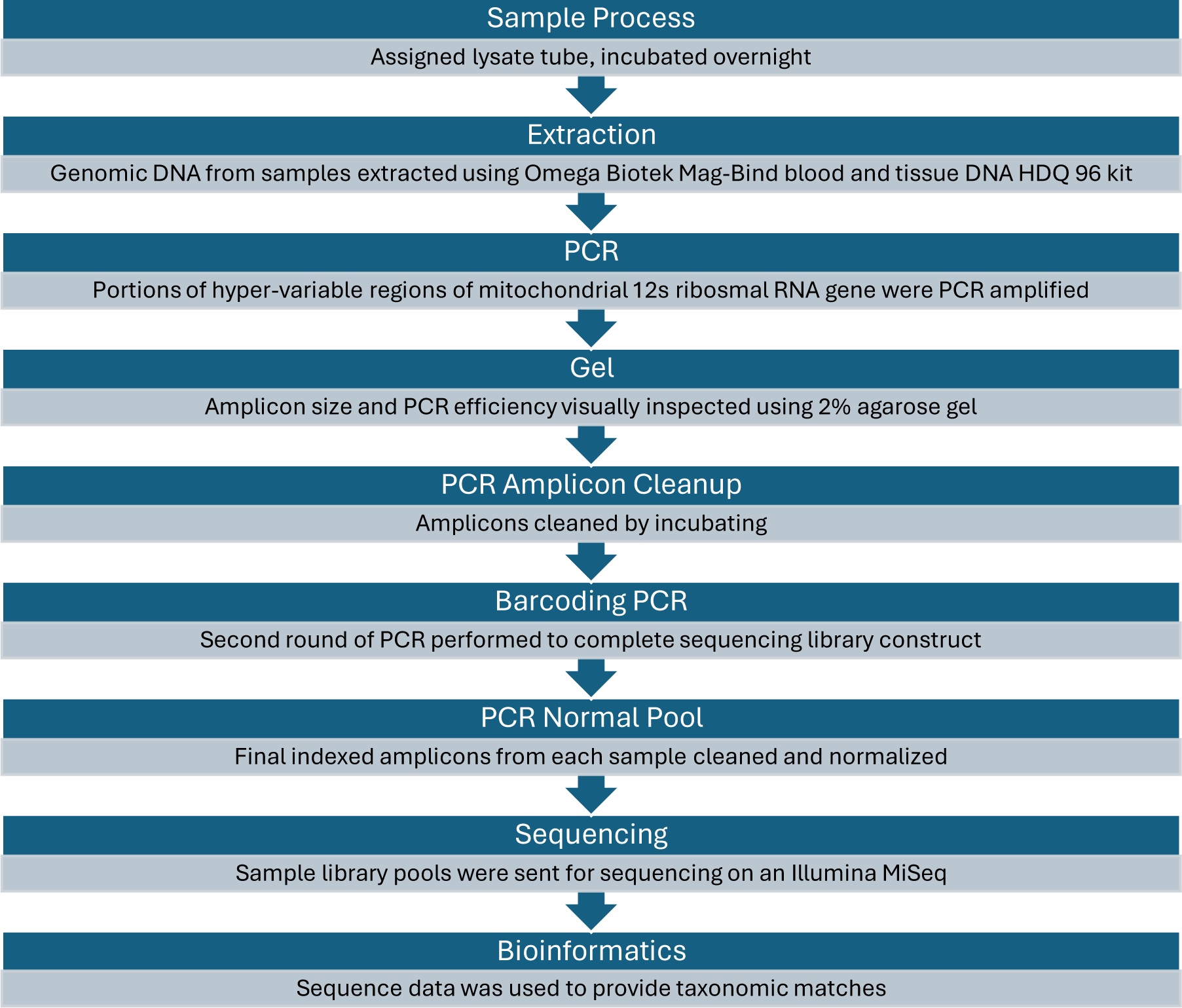
Flow chart of the 9-step process followed by Jonah Ventures to analyze eDNA samples.

The process used by Jonah Ventures was shared as follows. **Sample Process**, sample barcodes were recorded and assigned a corresponding lysate tube. Sample filters, lysis buffer, and proteinase K were heated to 56 C for one hour. Under a laminar flow hood, warm lysis buffers were pushed through the filter housing, and all supernatant was collected in the corresponding lysate tube. Tubes were placed in an incubator overnight at 56 C. After incubation the lysate tubes were immediately processed. **Extraction**, genomic DNA from samples was extracted using the Omega Biotek Mag-Bind Blood & Tissue DNA HDQ 96 Kit (4x96 Preps) (Cat. No. / ID: M6399-01) according to the manufacturer’s protocol. Whole (25mm or 47mm) filters were used for genomic DNA extraction. The extraction protocol was automated and completed using a Hamilton Microlab Starlet. Genomic DNA was eluted into 100 µl and frozen at - 20 C. **PCR**, Forward Primer: GTCGGTAAAACTCGTGCCAGC Reverse Primer: CATAGTGGGGTATCTAATCCCAGTTTG Primer reference: (Miya et al., 2015). Portions of hyper-variable regions of the mitochondrial 12S ribosomal RNA (rRNA) gene were PCR amplified from each genomic DNA sample using the MiFishUF and MiFishUR primers with spacer regions. Both forward and reverse primers also contained a 5’ adaptor sequence to allow for subsequent indexing and Illumina sequencing. PCR amplification was performed in replicates of six and all six replicates were not pooled and kept separate. Each 25 µL PCR reaction was mixed according to the Promega PCR Master Mix specifications (Promega catalog # M5133, Madison, WI) which included 12.5ul Master Mix, 0.5 µM of each primer, 1.0 µl of gDNA, and 10.5 µl DNase/RNase-free H2O. DNA was PCR amplified using the following conditions: initial denaturation at 95C for 3 minutes, followed by 45 cycles of 20 seconds at 98C, 30 seconds at 60C, and 30 seconds at 72C, and a final elongation at 72C for 10 minutes. **Gel**, to determine amplicon size and PCR efficiency, each reaction was visually inspected using a 2% agarose gel with 5µl of each sample as input. **PCR Amplicon Cleanup**, amplicons were then cleaned by incubating amplicons with Exo1/SAP for 30 minutes at 37C following by inactivation at 95C for 5 minutes and stored at -20C. **Barcoding PCR**, a second round of PCR was performed to complete the sequencing library construct, appending with the final Illumina sequencing adapters and integrating a sample-specific,12-nucleotide index sequence. The indexing PCR included Promega Master mix, 0.5 µM of each primer and 2 µl of template DNA (cleaned amplicon from the first PCR reaction) and consisted of an initial denaturation of 95 °C for 3 minutes followed by 8 cycles of 95 °C for 30 sec, 55 °C for 30 seconds and 72 °C for 30 seconds. **PCR Normal Pool**, final indexed amplicons from each sample were cleaned and normalized using SequalPrep Normalization Plates (Life Technologies, Carlsbad, CA). 25µl of PCR amplicon is purified and normalize using the Life Technologies SequalPrep Normalization kit (cat#A10510-01) according to the manufacturer’s protocol. Samples are then pooled together by adding 5µl of each normalized sample to the pool. **Sequencing**, sample library pools were sent for sequencing on an Illumina MiSeq (San Diego, CA) at the Texas A&M Agrilife Genomics and Bioinformatics Sequencing Core facility using the v2 500-cycle kit (cat# MS-102-2003). Necessary quality control measures were performed at the sequencing center prior to sequencing. **Bioinformatics**, raw sequence data were demultiplexed using pheniqs v2.1.0 [1], enforcing strict matching of sample barcode indices (i.e, no errors). Cutadapt v3.4 [2] was then used remove gene primers from the forward and reverse reads, discarding any read pairs where one or both primers (including a 6 bp, fully degenerate prefix) were not found at the expected location (5’) with an error rate < 0.15. Read pairs were then merged using vsearch v2.15.2 [3], discarding resulting sequences with a length of < 130 bp, > 210 bp, or with a maximum expected error rate [4] > 0.5 bp. For each sample, reads were then clustered using the unoise3 denoising algorithm [5] as implemented in vsearch, using an alpha value of 5 and discarding unique raw sequences observed less than 8 times. Counts of the resulting exact sequence variants (ESVs) were then compiled and putative chimeras were removed using the uchime3 algorithm, as implemented in vsearch. For each final ESV, a consensus taxonomy was assigned using a custom best-hits algorithm and a reference database consisting of publicly available sequences (GenBank [6]) as well as Jonah Ventures voucher sequences records. Reference database searching used an exhaustive semi-global pairwise alignment with vsearch, and match quality was quantified using a custom, query-centric approach, where the % match ignores terminal gaps in the target sequence, but not the query sequence. The consensus taxonomy was then generated using either all 100% matching reference sequences or all reference sequences within 1% of the top match, accepting the reference taxonomy for any taxonomic level with > 90% agreement across the top hits.

### R Visualization and Analysis

First, using detection data from both visits a Bayesian, single season, occupancy model, with flat priors with 3 MCMC chains and 10,000 draws from the Gibbs sampler, fitted in R version 3.6.1 using the wiqid package was made. This modeled 2 parameters psi (occupancy) and p (detection probability) of *Ambystoma*. Next, using our detections, pond sizes, temperature, and elevations collected we used these values in R version 3.6.1 to find if there is a correlation between the covariates and the binomial variable (if Tiger Salamanders were detected) 1 or 0. Detections of the tiger salamander is the discrete dependent variable while the covariates are independent variables. R version 3.6.1 was used during all steps of analysis. The packages used for analysis were: ggplot2, lme4, dplyr, moderndive, tidyverse, and wiqid. This was done with logistic regression due to the fact we are working with discrete data rather than continuous, logistic regression uses the glm function to fit a model to the data. All co-variates were modeled using logistic regression where the y axis is the binomial variable (response variable). The x axis was the numerical co-variate (predictor variable). Then the asymptote of the line was analyzed to find the point of interest. Top models were selected using AIC to show which model fit the data the best. Last the P-value of these models was used to determine if there was significance. When modeling elevation and surface area overall detection at the site was used. When modeling temperature for individual visits the specific detections and temperatures recorded during each visit were used, not over all detection at the site. Air temperature at visit 2 was the best model so one more model was ran with average temperature. This averaged the temperature at each site and used overall detection at each site to model. R scripts and corresponding csv raw data are posted to OSF open science framework account (https://osf.io/gkz8x/).

## Results

All 8 sample sites had filters stored from the visit on 6-24-23. But only 6 were collected on 7-29-23, because site 2 and site 8 lacked water. The highest elevation collected was at site 3 at 756.21 meters the lowest was site 5 at 688.85 meters (Figure 4). Temperature was collected for all sites except for site 2 and 8 on the second visit. The highest temperature recorded for first visit was 25° C at site 4 with the remaining sites all being 24° C (Figure 5). The highest during second visit was 26° C at sites 4 and 7, while the lowest was 22° C at site 1 (Figure 6). Surface area varied in size from smallest being site 7 at 40.74 meters squared and largest being site 5 at 14,609.15 meters squared (Figure 7).

**Figure 4.**
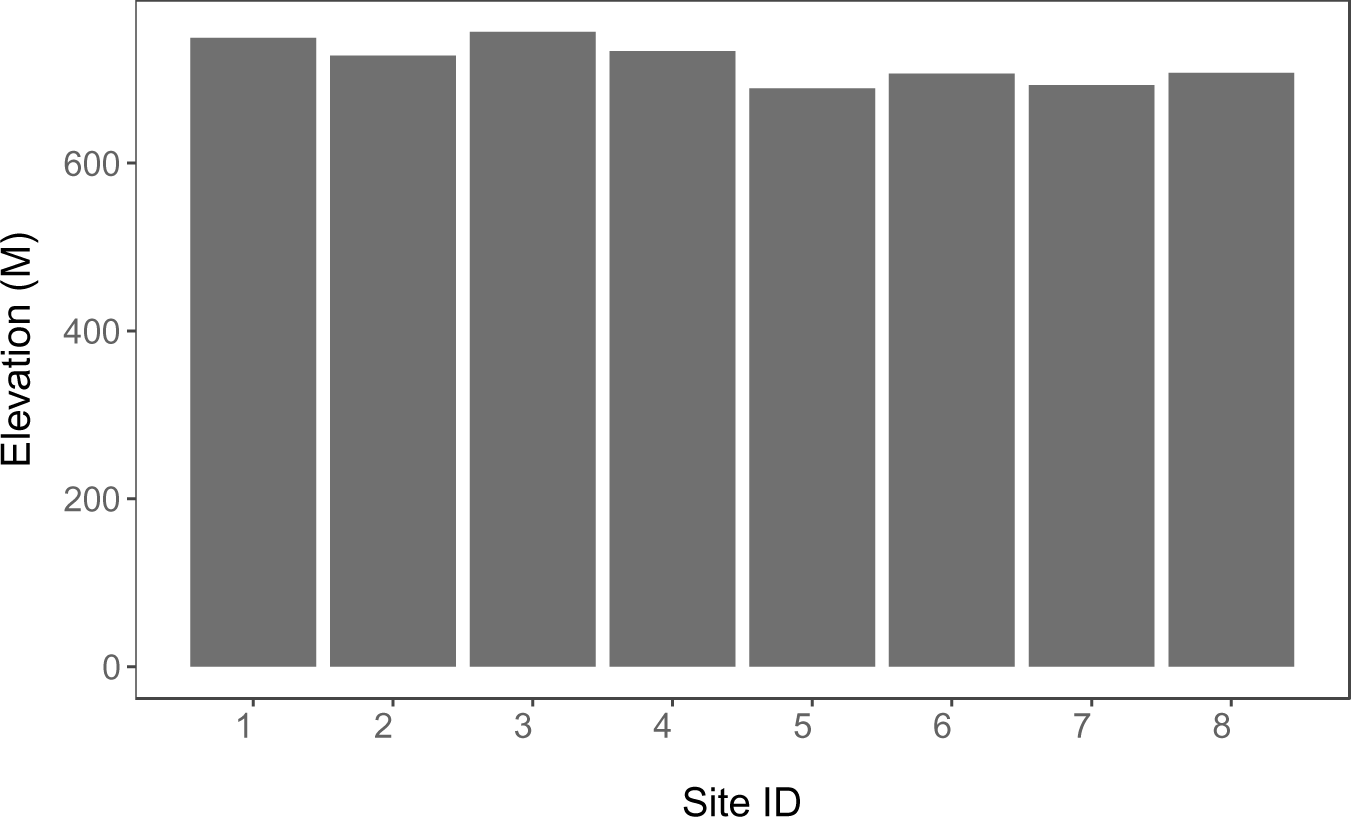
Bar chart displaying Site ID (x) and elevation in meters (y). Elevation data was obtained through GPS data provided by OnX maps, collected in feet then converted to meters. The figure shows a relatively uniform distribution of elevations across study sites. This model was produced using ggplot2 in R version 3.6.1.

**Figure 5.**
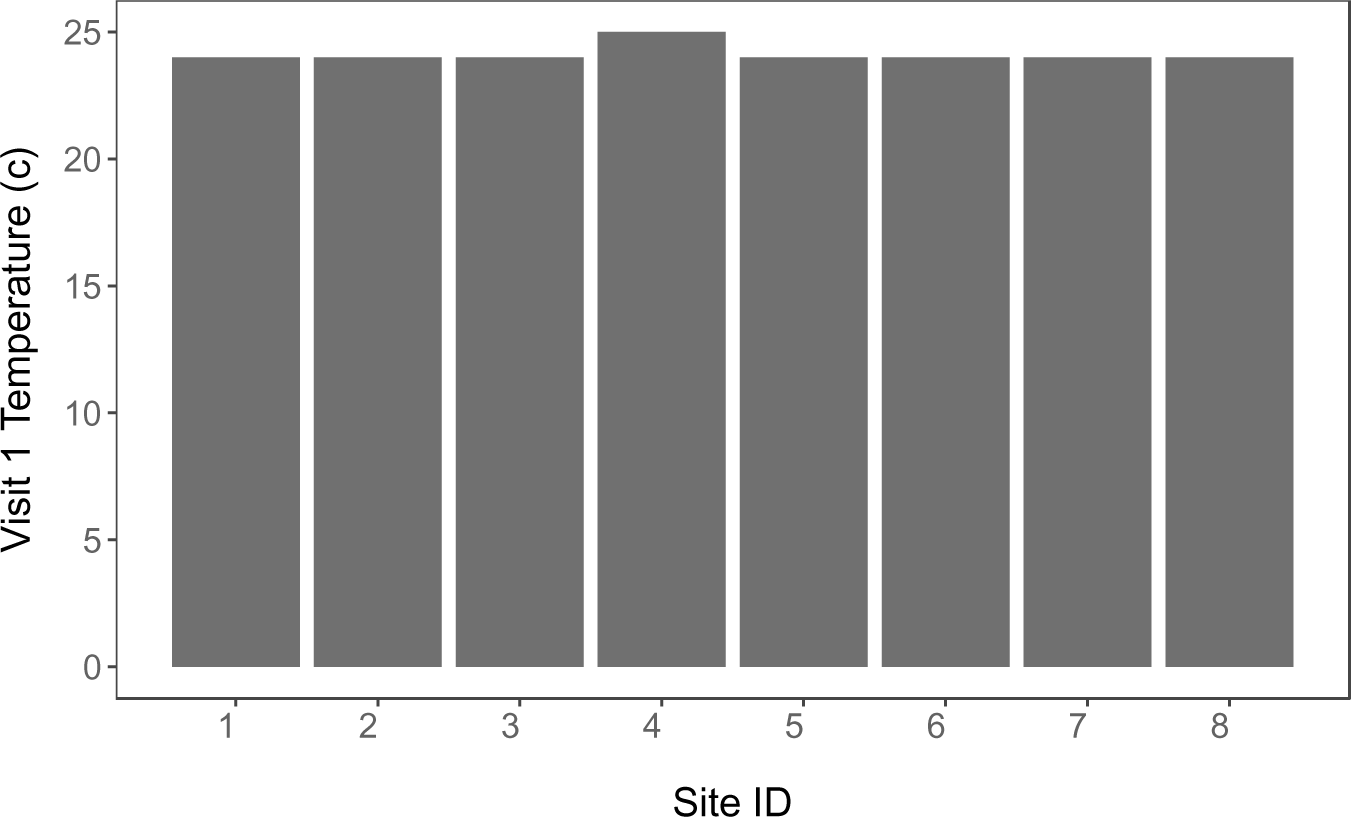
Bar chart displaying Site ID (x) and temperature on first visit in Celsius (y). Temperature data was obtained via iPhone data was collected in Fahrenheit then converted to Celsius. The figure shows a relatively uniform distribution of temperature across study sites. This model was produced using ggplot2 in R version 3.6.1.

**Figure 6.**
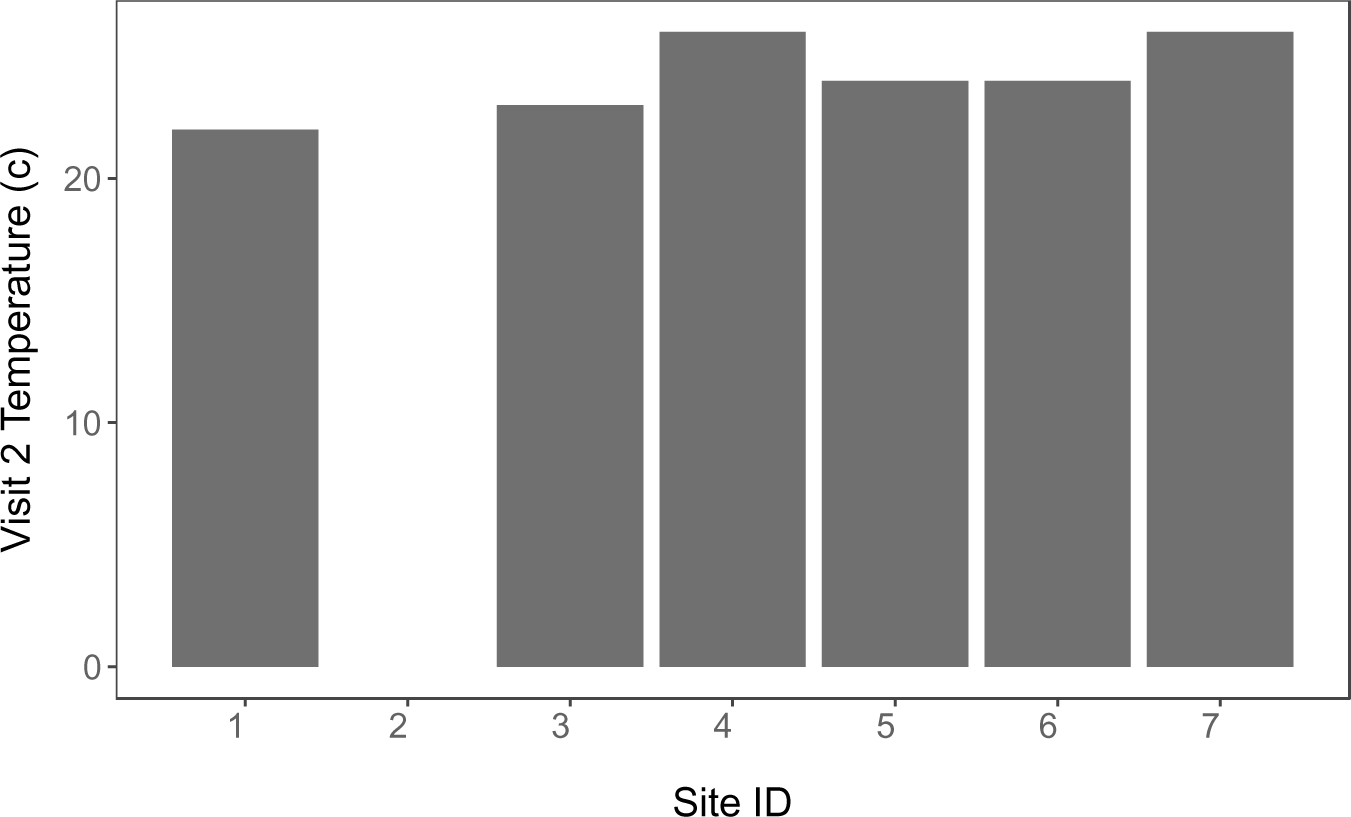
Bar chart displaying Site ID (x) and temperature on second visit in Celsius (y). Temperature data was obtained via iPhone data was collected in Fahrenheit then converted to Celsius. The figure shows a relatively uniform distribution of temperature across study sites. Sites 2 and 8 have no value as there were no data collected for these sites on the second visit due to the ponds being dry upon arrival. This model was produced using ggplot2 in R version 3.6.1.

**Figure 7.**
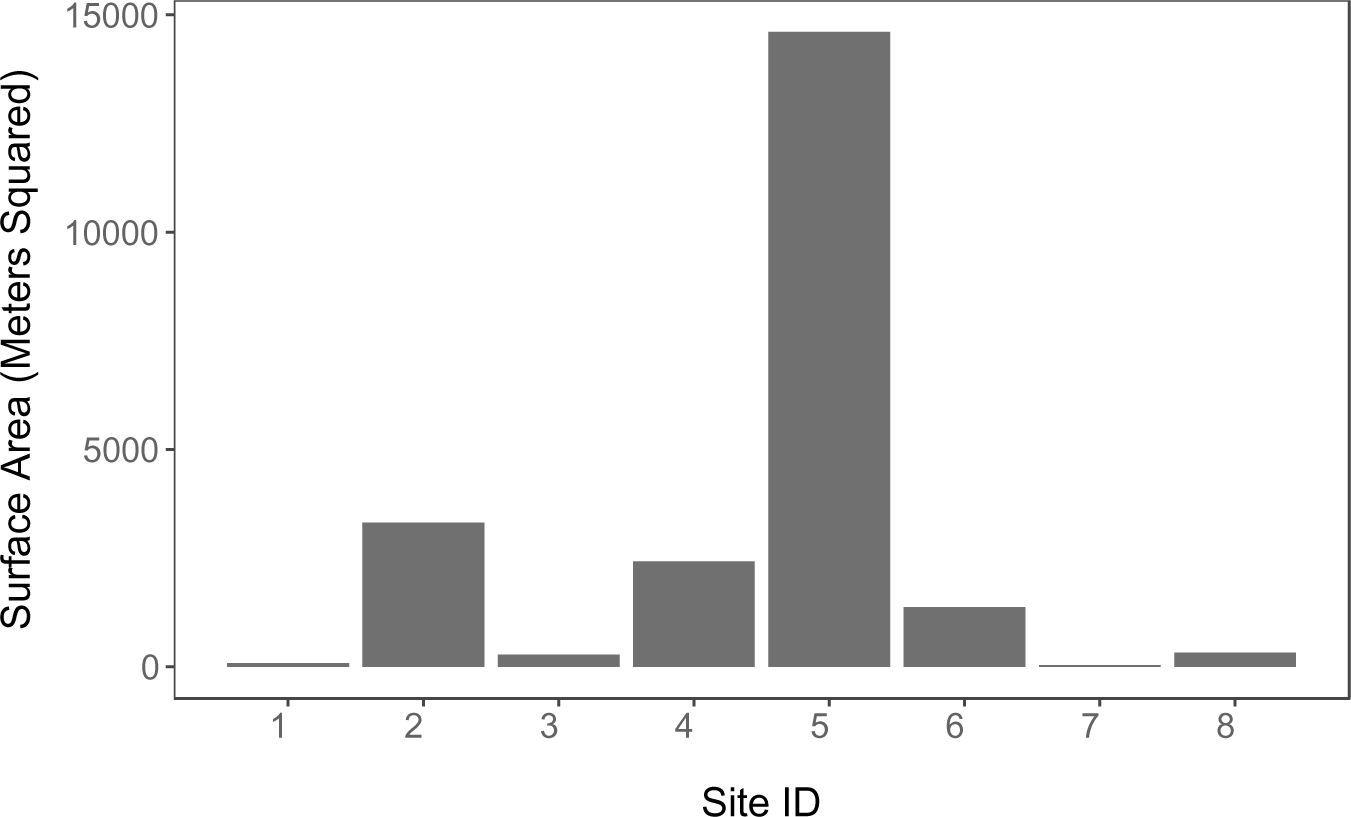
Bar chart displaying Site ID (x) and surface area in meters squared (y). Surface area data was obtained through OnX maps area measuring tool, collected in acres then converted to meters squared. The figure shows large amount of variance in surface area across study sites from 14,609.15 meters squared at site 5 and 40.74 meters squared at site 7. This model was produced using ggplot2 in R version 3.6.1.

There were positive detections for *ambystoma* at sites 1,2,4,6, and 8, and no detections at sites 3,5, and 7. Visit 1 had detections at sites 1,2,4,6, and 8. Visit 2 had detections at sites 4 and 6. Detection data was only certain down to the genus level of *Ambystoma* so that is what was modeled (Figure 8). We recevied 4 candidates for species level individuals in this genus all 1-5 base pairs away from being a 100% match. For psi (occupancy) we are 95% that 95% of our data is between .35 and .97 with a mean of .65. If we go to any pond in the study area there should be a 65% chance that it is occupied. For p (detection probability) we are 95% certain that 95% of our data is between .44 and .98 with a mean of .74. If we go to any pond in the study area that is occupied by *Ambystoma* there should be a 74% that we can detect them.

**Figure 8.**
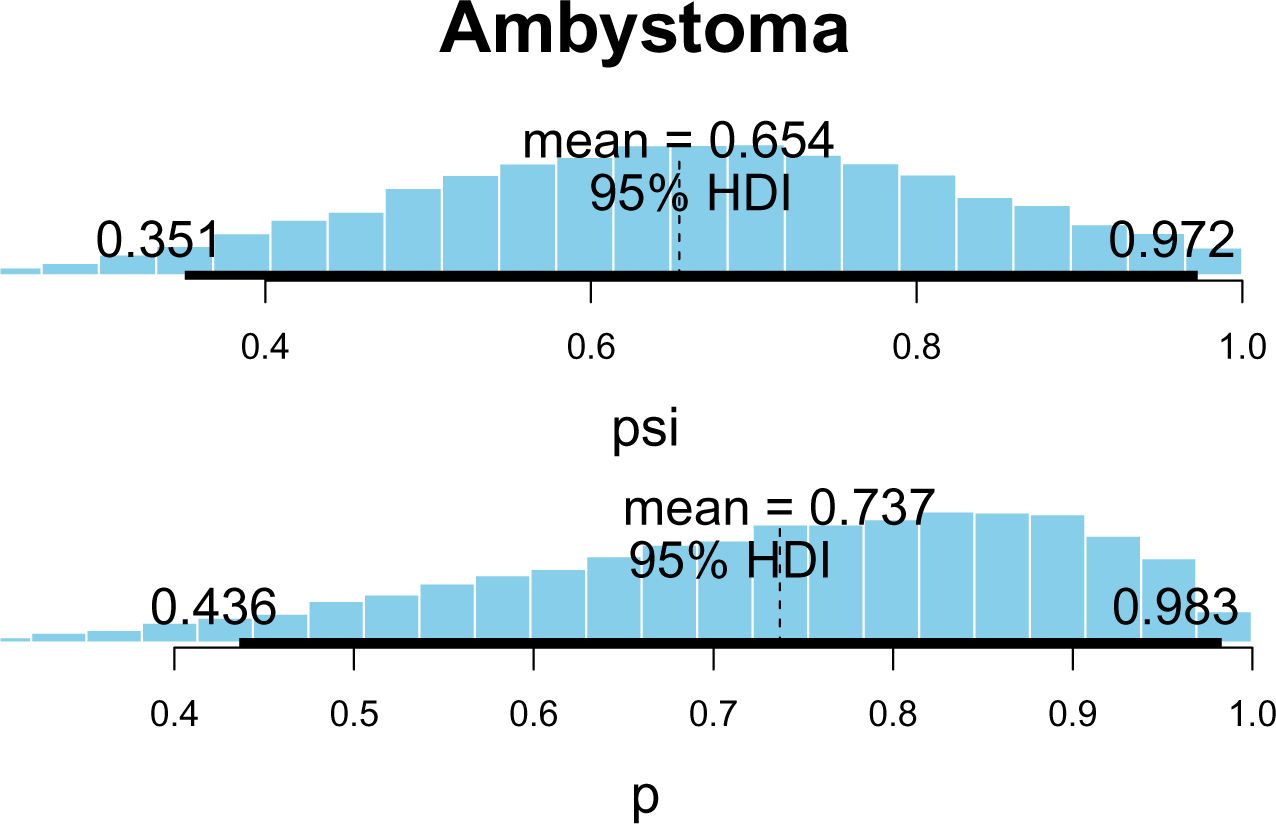
This is a Bayesian, single season, occupancy model, with flat priors with 3 MCMC chains and 10,000 draws from the Gibbs sampler, fitted in R version 3.6.1 using the wiqid package. It modeled 2 parameters psi (occupancy) and p (detection probability) of *Ambystoma*. Detection was obtained through metabarcoding of eDNA samples resulting in detection or not. This shows a mean occupancy of 65% and a mean detection probability of 74%.

Modeling detection against surface area shows us there is no strong correlation for detections and surface area having detections of yes and no from 40 meters squared up to 3500 with an outlier at 14000 (Figure 9). When detection was modeled against elevation there is no strong correlation having detections of yes and no ranging all the way across. But all our detections do come above 700 meters and below 750 meters (Figure 10).

**Figure 9.**
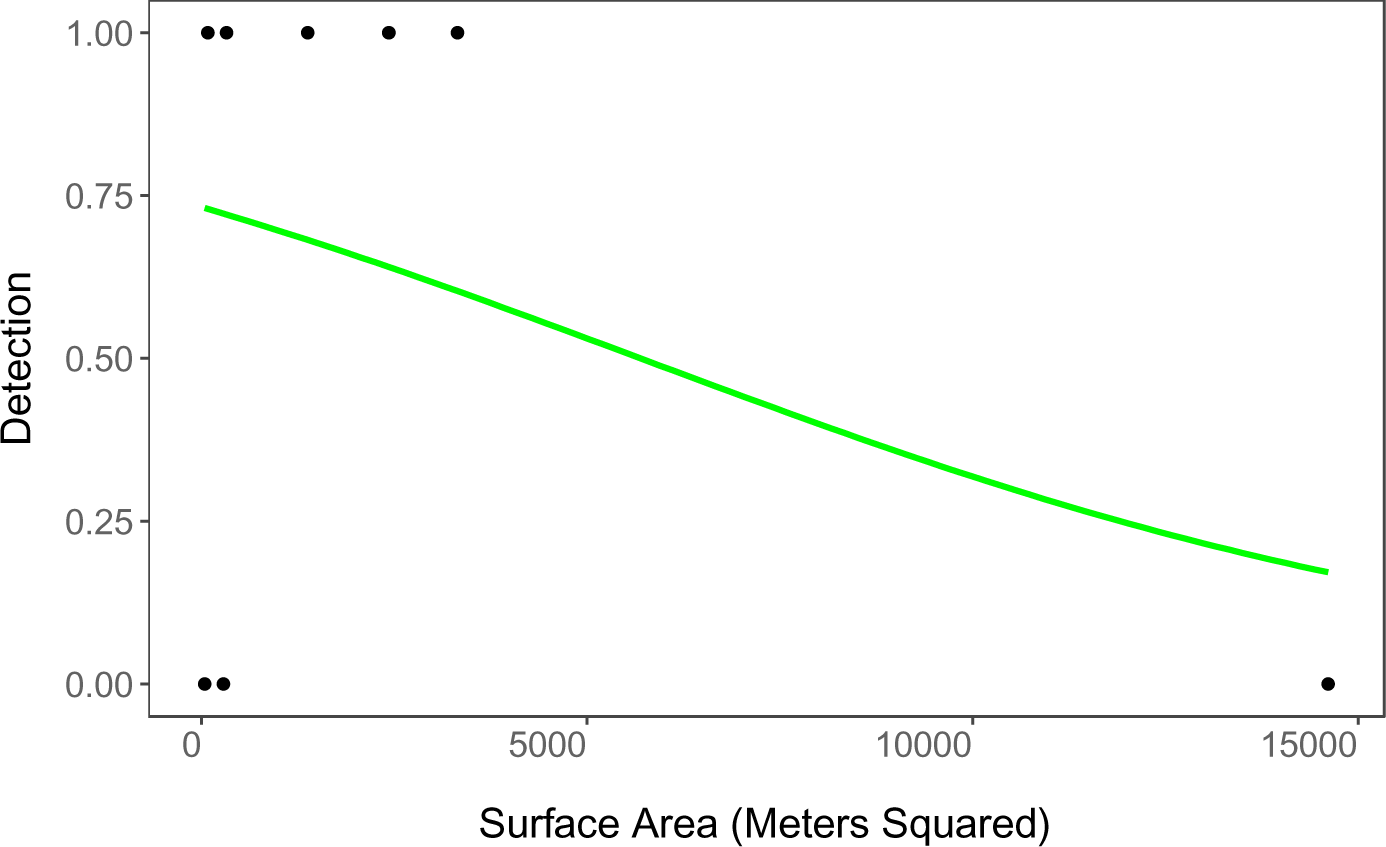
This is a generalized linear model using logistic regression displaying the relationship between the co-variate surface area in meters squared (x). As well as the binomial variable, detection indicated by 1 detected or 0 not (Y). Detection was obtained through metabarcoding of eDNA samples resulting in a detection or not. Surface area was measured with the area measuring tool on onX maps collected in acres and converted to meters squared. The figure shows no strong correlation for surface area and detections. This model was produced using ggplot2 in R version 3.6.1.

**Figure 10.**
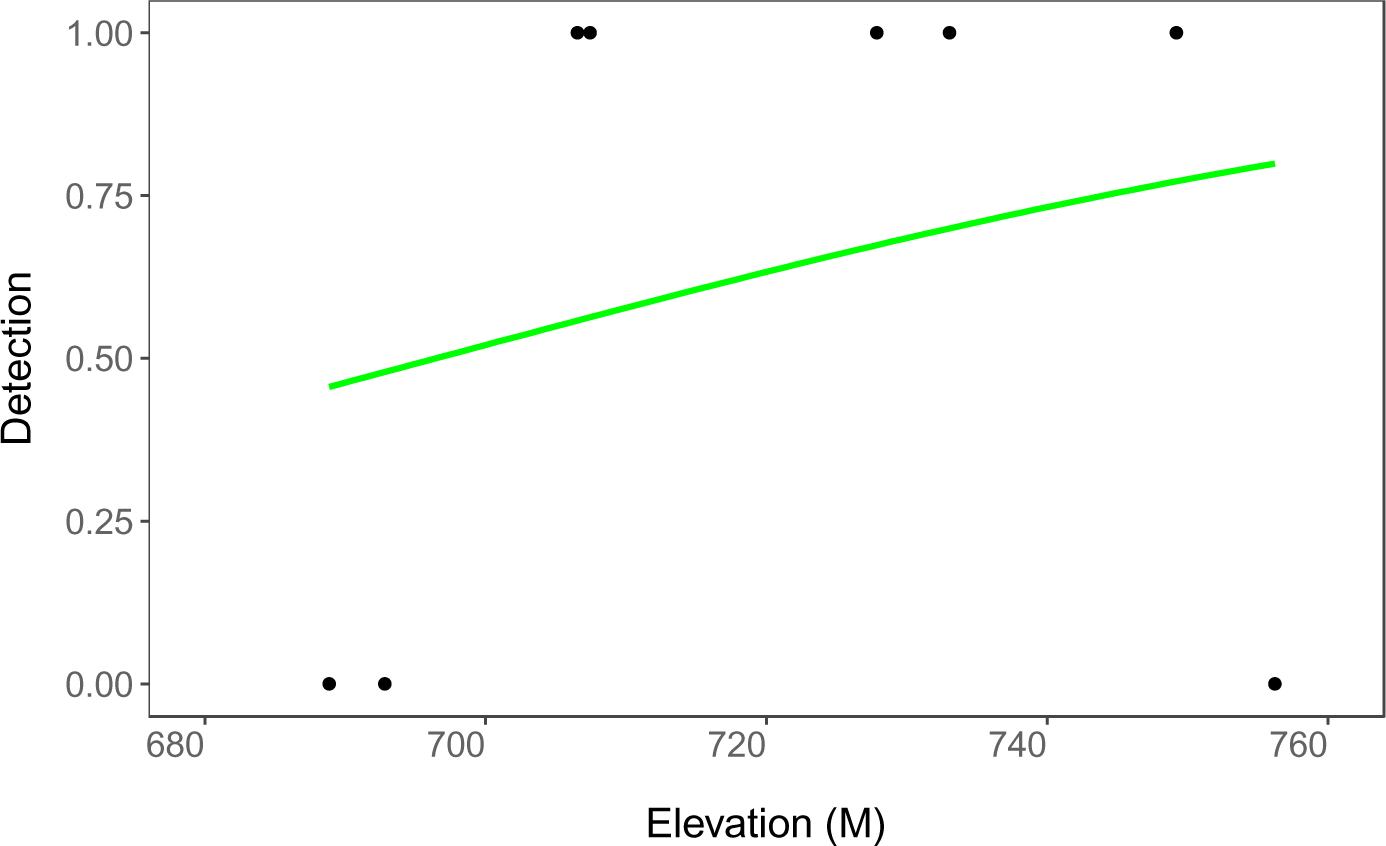
This is a generalized linear model using logistic regression displaying the relationship between the co-variate elevation in meters (x). As well as the binomial variable, detection indicated by 1 detected or 0 not (Y). Detection was obtained through qPCR of eDNA samples resulting in a detection or not. Elevation was measured with the GPS data on OnX maps. The figure shows no strong correlation for elevation and detections, but all detections happen above 700M and below 750M. This model was produced using ggplot2 in R version 3.6.1.

When temperature and detections for visit 1 were modeled there was not a strong correlation shown. All the temperatures recorded were at 24 C besides one at 25 C, there is not enough variation to see any major patterns with only 1 degree difference. (Figure 11). When modeling temperature and detections for visit 2 we can see a weak correlation where all the positive detections come at 24 C and after. With half of the no detections before 24 C and half at or after 24 C. (Figure 12). Average temperature shows no strong correlation with almost an even distribution of detections and no detections 3 and 3. The 3 detections and 3 no detections are almost in the same spot on the x axis as well, 23 C and 23.5 C, 24 C, 25 C and 25.5 C (Figure 13).

**Figure 11.**
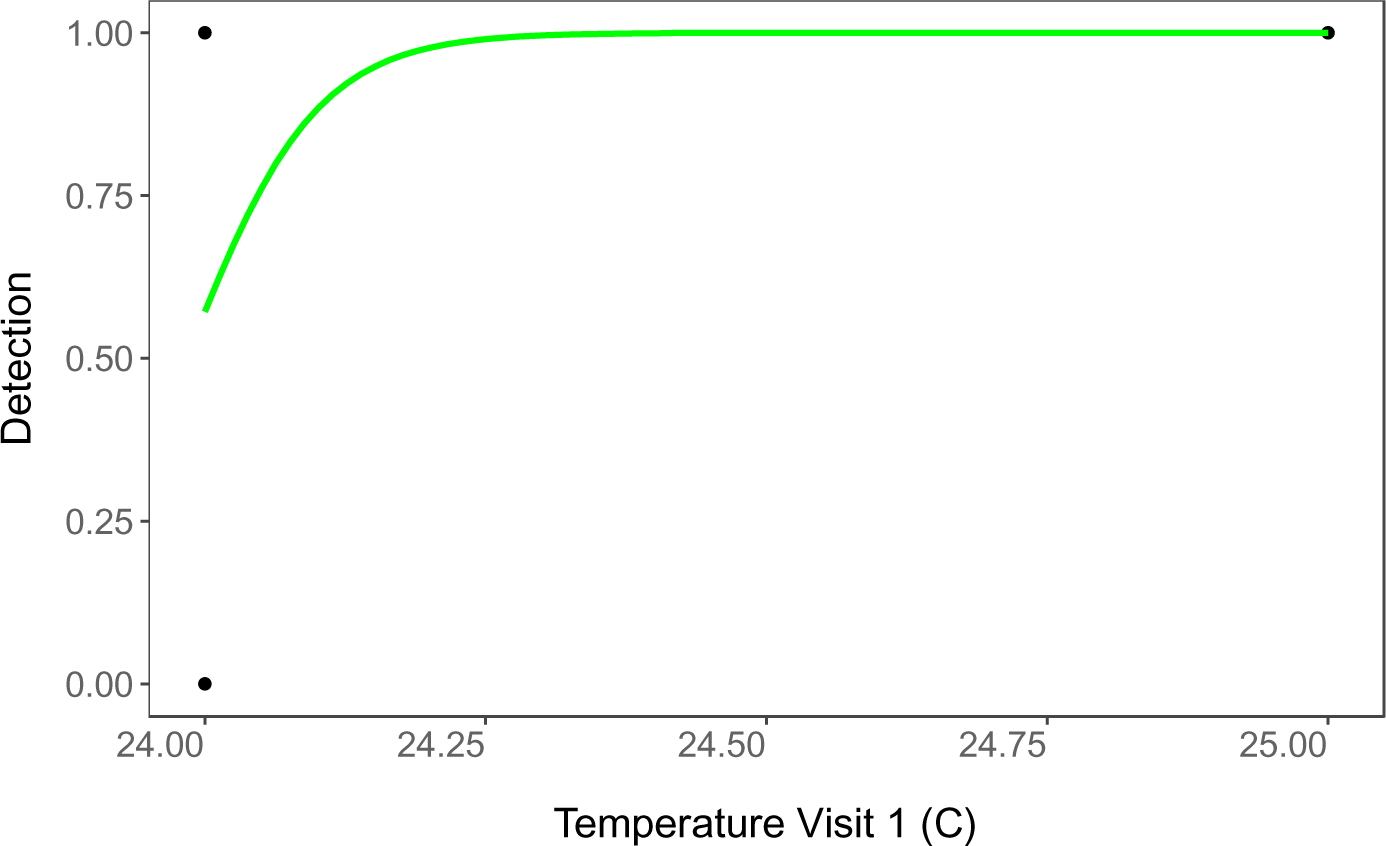
This is a generalized linear model using logistic regression displaying the relationship between the co-variate temperature on visit 1 in Celsius (x). As well as the binomial variable, detection indicated by 1 detected or 0 not (Y). This was from visit 1 on 6-24-23. Detection was obtained through qPCR of eDNA samples resulting in a detection or not. Temperature was measured with weather app on iPhone at time of sample collection collected in Fahrenheit and converted to Celsius. The figure shows no correlation between Detections and Temperature at visit 1. Not enough variation to see trends with all points being at 24 C and 25 C. This model was produced using ggplot2 in R version 3.6.1.

**Figure 12.**
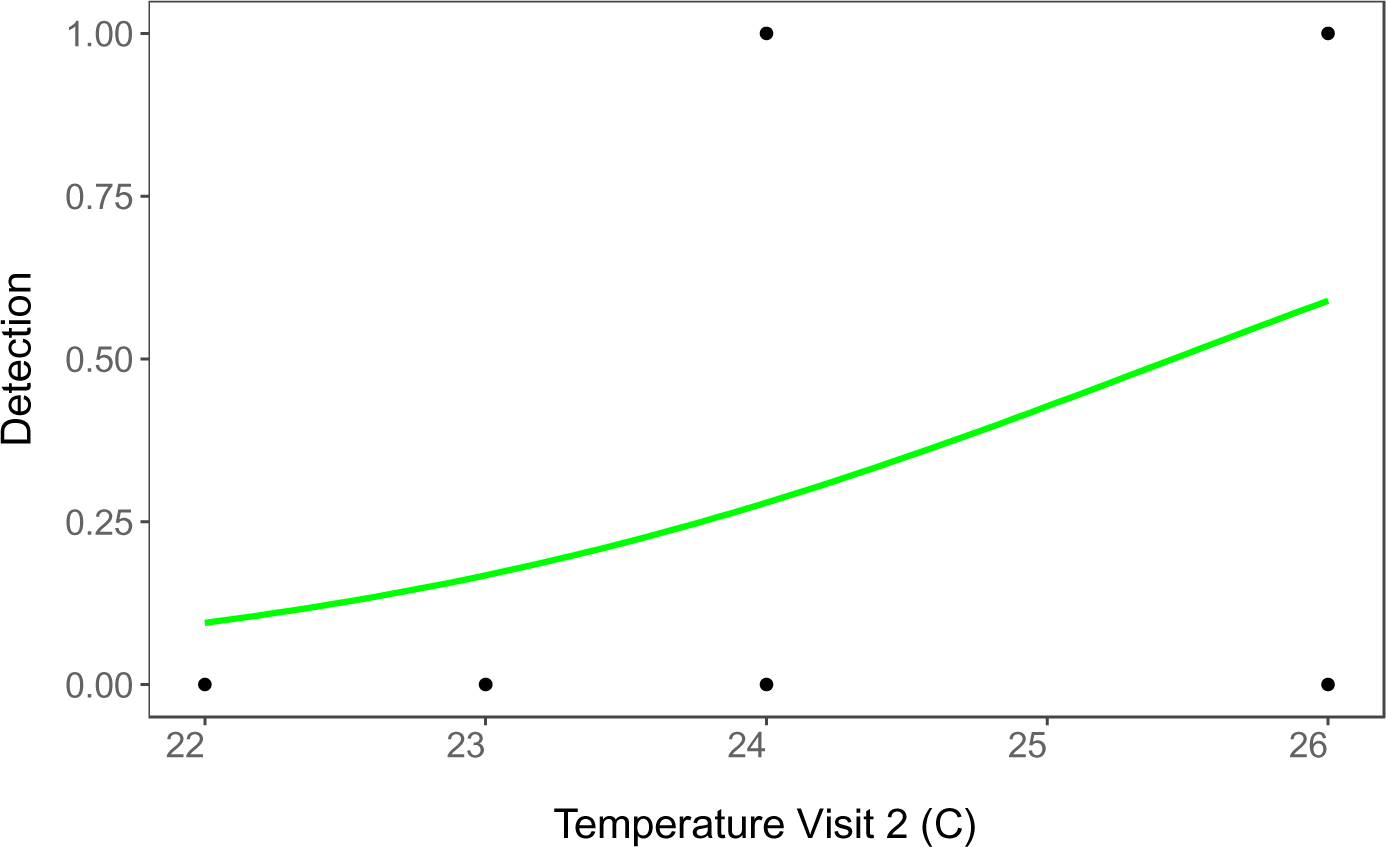
This is a generalized linear model using logistic regression displaying the relationship between the co-variate temperature on visit 2 in Celsius (x). As well as the binomial variable, detection indicated by 1 detected or 0 not (Y). This was from visit 2 on 7-29-23. Detection was obtained through qPCR of eDNA samples resulting in a detection or not. Temperature was measured with weather app on iPhone at time of sample collection collected in Fahrenheit and converted to Celsius. The figure shows a weak correlation where all detections come at or after 24 C. While half of the no detections are before 24 C and half are at or after 24 C. This model was produced using ggplot2 in R version 3.6.1.

**Figure 13.**
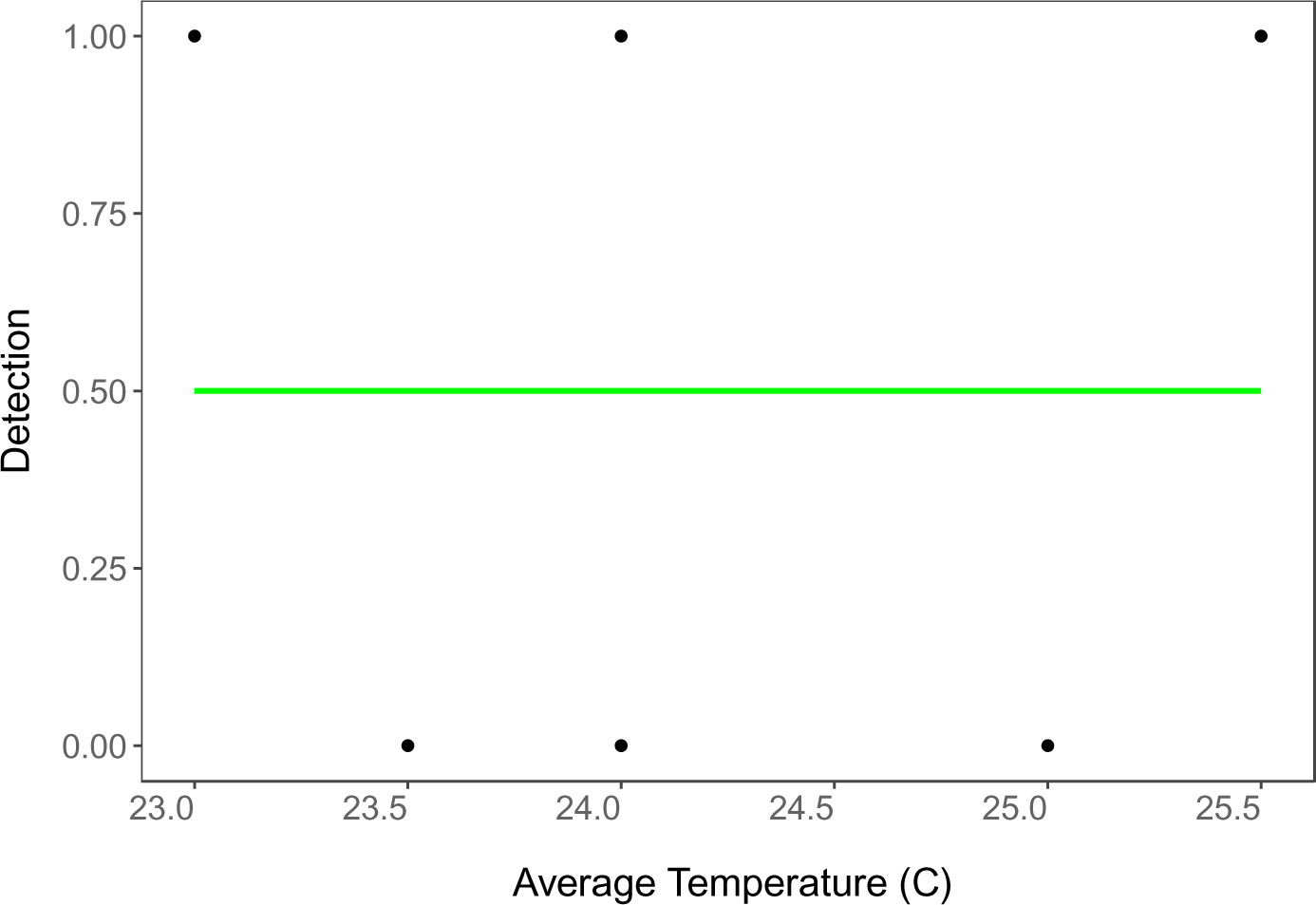
This is a generalized linear model using logistic regression displaying the relationship between the co-variate average temperature at each site in Celsius (x). As well as the binomial variable, detection indicated by 1 detected or 0 not (Y). This was average temperature for each site from visit 1 on 6-24-23 and visit 2 on 7-29-23. Detection was obtained through qPCR of eDNA samples resulting in a detection or not. Temperature was measured with weather app on iPhone at time of sample collection collected in Fahrenheit and converted to Celsius then combined and averaged for each site. The figure shows no strong correlation with a almost uniform distribution between detections and no detections. This model was produced using ggplot2 in R version 3.6.1.

When comparing our 5 models based on AIC value the model that fits the data the best is air temperature at visit 2 with an AIC value of 10.61. This is a 3.46 difference from elevation which based on AIC value fits the data the worst. Average temperature is the second-best model with an AIC value of 12.3, a 1.7 difference from our best model. Surface area and air temperature at visit 1 are very similar in AIC value and have the same model likelihood and model weight. Although air temperature at visit 2 is the model that fits the data the best, based on its P-value of .35 it does not satisfy our alpha critical value of .01 so it is not significant. (Table 1).

**Table 1.** This an AIC table displaying degrees freedom, AIC value, Delta AIC, Model Likelihood, and Model weight. For the 5 collected models, air temperature at visit 1 (AT_v1_c), air temperature at visit 2 (AT_v2_c), average air temperature at each site from both visits (average_temp), elevation, and surface area (meters). The model shows the same degrees freedom with the best fitting model being air temperature at visit 2 with the worst fitting being elevation with a difference of 3.46. Air temperature at visit 2 is the best fitting model but still not significant based on P-value. This model was produced using wiqid in R version 3.6.1.

| Model Parameters | df | AIC | $\Delta$ AIC | Model Likelihood | Model Weight |
| --- | --- | --- | --- | --- | --- |
| Detection~AT_v2_c | 2 | 10.61 | 0 | 1 | 0.48 |
| Detection~average_temp | 2 | 12.3 | 1.70 | 0.43 | 0.21 |
| Detection~meters | 2 | 13.52 | 2.91 | 0.23 | 0.11 |
| Detection~AT_v1_c | 2 | 13.56 | 2.95 | 0.23 | 0.11 |
| Detection~elevation | 2 | 14.07 | 3.46 | 0.18 | 0.09 |

## Discussion

We were able to detect *Ambystoma* at 5 of the 8 randomly selected sites. The genus *Ambystoma* is the highest taxonomic level that was able to be accurately detected. When detection data was analyzed using Bayesian single season occupancy analysis the mean psi (occupancy) was .65 and the mean p (detection probability) was .74. Upon running logistic regression to model the covariates it was found that air temperature at visit 2 was the model that fit the data the best. However, given the P-value of .35 for this model it was not significant. None of the covariates were able to give insight into what affects if detections were possible at each site. The occupancy analysis provided promising results however. Providing support for the efficiency of PCR and its ability to detect cryptic species. Going forward the goal would be to answer the question of what affects the detection of *Ambystoma.* Finding covariates that have a correlation to detections.

The original goal was to detect the species *Ambystoma tigrinum* (tiger salamander) as that is the species of salamander present in this area. We were successful in getting detections in 5 of the 8 randomly selected sites. However, the detections could not be backed by data all the way down to a species level. We were only able to be confident up to the genus level of *Ambystoma.* Upon receiving the results from the lab we had 4 possible candidate species; *Ambystoma mexicanum* (Axolotl)*, Ambystoma mavortium* (Barred tiger salamander)*, Ambystoma tigrinum* (tiger salamander), and *Ambystoma andersoni* (Andersons salamander). The axolotl was a 99.5% match, barred tiger salamander was a 98.9% match, tiger salamander was a 98.9% match, and the Andersons salamander was a 98.9% match. This means that these individuals were between 1-5 base pairs off being a 100% match. Combined with data from the North Dakota Game and Fish it could be hypothesized that the species found was the tiger salamander as the others are not reported in the study area. However, off data strictly we can only be confident to a genus level.

There are some basic observations that are interesting to note about our detection data. The first of which is the 2 sites that lacked water upon arrival for the second round of sampling both yielded positive detections in the first round of sampling. This is interesting to note, however in amphibians it is common to reproduce in areas known as vernal pools which are season pools of water (Semlitsch & Skelly, 2008). The other interesting thing to note is the 3 sites which did not yield a detection ever (sites 3,5,7). Site 3 was unique in that it had some water running into it which was different from all the other sites. Site 5 was the largest of all sites collected from in terms of surface area in meters squared at 14,609.15. Lastly, site 7 was not measured for iron content however, it did have a natural spring running into it and it was a very dark reddish brown colored water. If this was high iron content it would make sense no salamanders were living there being they are a sensitive species because they permeate through their skin.

Running a Bayesian single season occupancy analysis in this study allowed us to examine the occupancy and detection probability of *Ambystoma* in this area using an eDNA collection method. When doing this study we were able to collect samples from all sites in one 6 hour day. It cost $105 per sample and we were able to get a read out of every species that was detected in each sample with percent match of taxonomy not just the target species. The analysis tells us that if we were to go to any pond in the study area there should be a 65% chance that the pond is occupied by *Ambystoma.* It also tells us that if we were to go to any pond in the study area that is currently occupied by *Ambystoma* there should be a 74% chance that we detect them. This is a very efficient detection rate and adds support to the efficiency of eDNA in detecting cryptic species. No other sampling methods were employed to compare to the eDNA collection method to. Although, in a future study this would be an elucidating piece of the puzzle comparing eDNA to more traditional methods. (Andres et al., 2023) examined this comparing eDNA to traditional methods including electrofishing, gillnetting, fyke netting, and seining. They found that eDNA found many more unique species than any other method. 10 more species than electrofishing, 26 more than gillnetting, 14 more than fyke netting, and 14 more than seining.

Logistic regression modeling of our covariates provided no correlations or answers. The model that fit the data the best based off the lowest AIC value was air temperature at visit 2 with an AIC of 10.61. Our alpha critical value for this study was set at 0.01, requiring a P-value lower than this to yield significance. The P-value for air temperature at visit two was .35. Meaning even our best fit model does not have any significance. In future studies connecting this piece of the puzzle would be very interesting in determining a covariate that can have correlation to detections. (Strickler, Fremier & Goldberg, 2015) examine eDNA degradation and found that uv-B radiation, temperature, and pH level affect the degradation rate of eDNA. Indicating that a lower temperature, more basic pH, and protection from uv-B radiation can help with the degradation of eDNA. Moving forward this is a place to start to expand upon this work.

## Conclusion

This eDNA sampling method was successful in obtaining positive detections of *Ambystoma* at 5 of the 8 total sites. Bayesian single season occupancy analysis provided an occupancy of 65% and a detection probability of 74% at these sites. Providing support for eDNA as an efficient method in detecting cryptic species. The covariates collected provided no significance and thus no correlations to detections. This study was successful in providing support for the efficiency of eDNA sampling in the detection of cryptic species however it was unsuccessful in finding significant covariates.

## Acknowledgments

Thank you to Dr. Colin Strine for help in analysis and overall guidance on this project. Trevor Hann M.S. for guidance and help in analyzing field samples. Jonah Ventures for analyzing eDNA samples through metabarcoding. National Genomics Center for Wildlife and Fish Conservation, as well as the Rocky Mountain Research station for all sampling equipment and sampling protocol. Kreidt Ranch for land access and introduction to other landowners as well as help in sampling.

